# Sprague-Dawley rats differ in responses to medial perforant path paired pulse and tetanic activation as a function of sex and age

**DOI:** 10.1101/2022.10.05.511027

**Authors:** Susan G. Walling, Carolyn W. Harley, Gerard M Martin, Olivia DE Dutton, Alexander T Burke, Ella A Chirinos

## Abstract

Network plasticity in the medial perforant path (MPP) of adult (5-9 mo) and aged (18-20 mo) urethane-anesthetized male and female Sprague-Dawley rats was characterized. Paired pulses probed recurrent networks before and after a moderate tetanic protocol.

Adult females exhibited greater EPSP-spike coupling suggesting greater intrinsic excitability than adult males. Aged rats did not differ in EPSP-spike coupling but aged females had larger spikes at high currents than males. Paired pulses suggested lower GABA-B inhibition in females.

Absolute population spike measures were larger post-tetani in female rats than male rats. Relative population spike increases were greatest in adult males relative to females and to aged males. EPSP slope potentiation was detected with normalization in some post-tetanic intervals for all groups except aged males. Tetani shortened spike latency across groups.

Tetani-associated NMDA-mediated burst depolarizations were larger for the first two trains in each tetanus in adult males than other groups. EPSP slopes over 30 min post-tetani predicted spike size in female rats, but not in males.

Replicating newer evidence MPP plasticity in adult males was mediated by increased intrinsic excitability. Female MPP plasticity was related to synaptic drive increases, not excitability increases. Aged male rats were deficient in MPP plasticity.

## INTRODUCTION

In 1973, Bliss and Lomo published a landmark paper demonstrating tetanus-induced long-term potentiation (LTP) in medial perforant path (MPP)-evoked potentials recorded in the dentate gyrus of anesthetized rabbit^1^. This paper became the starting point for hypotheses concerning the physiological basis of learning and memory. The enduring plasticity Bliss and Lomo found most frequently was reduced population spike (PS) latency, but increases in population excitatory post-synaptic potentials (EPSPs) generated the greatest interest. Increases in PS amplitude were similarly frequent, but not consistently correlated with EPSP increases. Bliss and Lomo conclude that two independent mechanisms were responsible for MPP-LTP: (a) an increase in the efficacy of synaptic transmission and (b) an increase in the excitability of the granule cell population. The preponderance of experimental LTP investigations in dentate gyrus focused on the (a) mechanism, EPSP potentiation. However, in the last two decades, attention has turned to excitability increases. In 2016, Lopez-Rojas et al.^2^ presented evidence that an increase in dendritic intrinsic excitability is primarily responsible for dentate gyrus MPP-LTP in mature granule cells of male rats. A comparative review of EPSP and intrinsic excitability changes in learning and memory^3^ highlights the commonality of their induction mechanisms with both depending on NMDA receptors. Further, both types of plasticity events may act bi-directionally, increasing or decreasing connectivity, in neural networks. Sex and age differences have been understudied in MPP plasticity. The present set of experiments address those variables in the context of Bliss and Lomo’s original observations.

## MATERIALS AND METHODS

Male and female Sprague Dawley rats (Charles River, QC) were housed doubly in individually ventilated cages (Techniplast, CA) with regular enrichment, on a reversed light cycle 12h:12h (lights off at 07.00h) until the age of ∼2-4mo, and then singly housed thereafter. Rats were fed regular chow (Teklad2018) however, both male and female rats were placed on a modest food restriction schedule at 2-3mo of age, to maintain a healthy aging weight profile and to reduce obesity related illnesses in old age ^4^. Rats were fed between 08.30h-10.30h daily and received an amount of food that was 75% (g) of the average age- and sex-dependent ad libitum consumption^4^. Water was available ad libitum. The average mass at the time of electrophysiological recording in the 5-7mo old rats; females 366.33g±27.45 and males 674.8g± 38.89, and for 18-20mo old rats; females 456.00g±81.82 and males, 748.80g±71.11.

All experimental procedures occurred within the dark phase of the animals’ light cycle (09.00h – 17.00h) and were performed in accordance with the Canadian Council of Animal Care (CCAC) guidelines and approved by the Memorial University of Newfoundland Institutional Animal Care Committee.

### Electrophysiological Recording

At 5-9mo or 18-20mo of age, rats were anesthetized with urethane (i.p.). To reduce overdose susceptibility due to age, sex or food restriction^5^, rats were stepped up to a ∼1.5g/kg dose (15% w/v). Once anesthetized the rats were placed in a stereotaxic instrument in the skull flat position and body temperature was maintained at 37±0.5°C via a feedback-regulated heating pad (FHC; ME). A concentric bipolar stimulating electrode (NE-100; Kopf, CA, USA) was lowered into the PP (∼7.2mm posterior, and ∼4.1 lateral to bregma, and 3.0mm ventral from brain surface) and a glass pipette (0.9% NaCl, 1-3MΩ) was lowered into the DG (∼3.5 mm posterior and ∼2.0mm lateral, and 2.5mm ventral; adjusted for animal size). A stainless-steel jeweler’s screw (Fine Science Tools, CA) served as ground. A 0.2ms square unipolar test pulse was delivered (0.1 Hz) to the PP and the DG responses were filtered 1-Hz – 10kHz, digitized at 10kHz and stored online using SciWorks 9.0 or 11.0 software (Datawave Technologies, CO). The stimulating and recording electrodes were then adjusted in the dorsal/ventral plane so that a maximal positive going waveform (granule cell layer) was achieved. See graphical experimental procedures in Supplemental Material Figure 1 (S1) for outline of experimental procedures and analyses.

**Figure 1.**
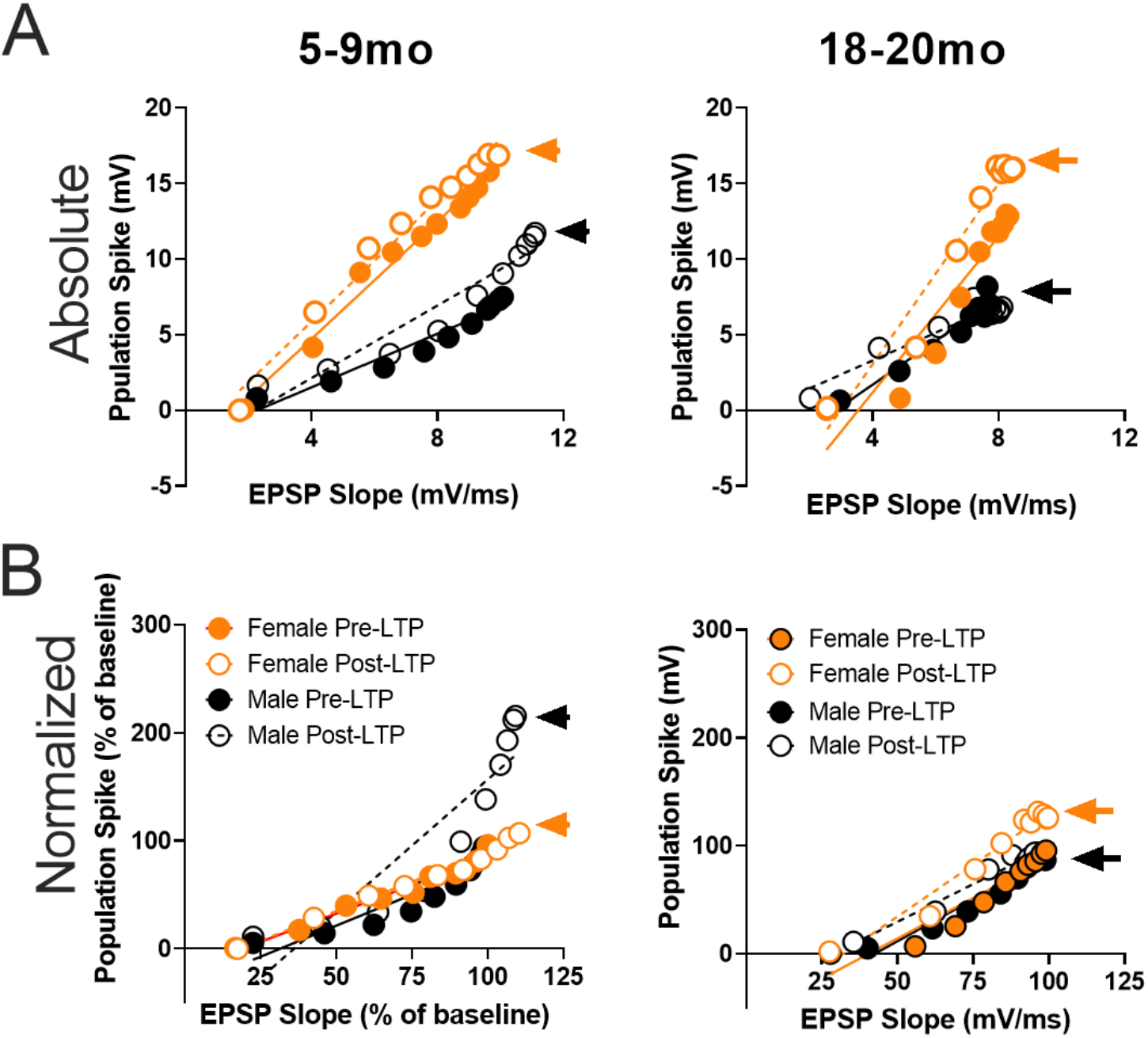
The effects of a moderate strength tetanic LTP protocol on E-S coupling of the medial perforant path input to dentate gyrus E-S coupling from I-O pre- and post-LTP curves in adult and aged adult, male and female rats. **A**. Absolute EPSP and PS values in adult (5-9mo) rats indicate a leftward shift in E-S coupling values in female rats, and values indicative of a ceiling effect post-LTP (orange dashed arrow). In aged male rats (18-20mo) PS values are approximately 50% of female PS values (solid arrows). **B**. Relative E-S plot with data normalized to the largest average PS and EPSP slope values from the Pre-LTP I-O curve, illustrate that aged female rats express increased PS increases for similar pre- and post-LTP EPSP slope values, suggesting higher levels of intrinsic excitability support post-tetanic potentiation of the MPP input to DG (solid orange arrow).

### Current Intensity E-S Coupling and Paired Pulse Analyses

At the commencement of the experiment, an input-output (I-O) current intensity curve (100-1000µA, 100µA steps) was performed using a series of three paired pulses presented every 10sec at each current level. Sets of paired pulses consisted of one presentation each of interstimulus intervals (ISI) 15ms, 70ms, and 120ms such that each current level tested each ISI before the current was increased. The paired pulse intervals were chosen to probe GABA-A sensitive paired pulse inhibition (PPI, 15ms), paired pulse facilitation (PPF, 70ms) and GABA-B late paired pulse inhibition (PPI, 120ms) of the PP-DG evoked PS in urethane anesthetized male^6^and female^7^ rats. The current level used for baseline stimulation during the PP-DG LTP experiment was the current producing a PS on the first (P1) of the two evoked paired responses that was ∼50% of maximum PS amplitude achieved in the I-O current curve. These current-paired pulse procedures were performed again at the termination of the LTP recording procedures to determine the effect of LTP on current intensity analysis and paired pulse facilitation and inhibition.

I-O current E-S coupling analysis was performed similarly to Walling and Harley^8^. In brief, each of the average of the three PS and EPSP slope measurements at each current level in the pre- and post-LTP curves, were contrasted across a range of EPSP measures (low current, smaller EPSP slope; through higher current and larger measures of the synaptic response). Measures are presented as absolute values (mV/ms and mV for slope and PS, respectively) and normalized to the largest, average PS and EPSP slope from the pre-LTP current curve.

Paired pulse ratios (PPR, pulse2/pulse1; P2/P1) were calculated for the PS amplitude at each ISI, and at each current intensity level. For presentation, the P2/P1 ratio from the stimulation intensity used for the LTP experiment (baseline; ∼50% maximal PS) was contrasted with P2/P1 ratio at the same current intensity after LTP (post-LTP). For PPR current intensity analyses (Supplemental Material Figure S4) outlier PPR values, in which the population spike amplitude on the second pulse was >3 standard deviations (s.d.) above normal, PPR values were capped to +2 s.d. of the highest PPR value for the animal at the respective ISIs. This treatment was applied to 4/360 PPR values in the 6-8mo old male rats and 3/360 values in 18-20mo old female rats.

### Moderate Strength Burst Long-Term Potentiation (LTP) Stimulation

After completion of the initial I-O and paired pulse curves, a tetanic LTP protocol was tested. This protocol of the three tested by Straube and Frey^9^, the moderate Protocol B, produced robust, β-adrenergic receptor, and protein synthesis-dependent, LTP of the PS variable in behaving male Wistar rats. The PP was stimulated at 0.1Hz with a monophasic 0.2ms pulse and after 30min of stable baseline evoked responses, monophasic tetanic stimulation at 200Hz was applied, consisting of 20 trains of 15 pulses (0.25ms width) with 10sec interburst intervals. Following the tetanic stimulation, the PP-DG evoked responses were followed for 90min recording followed by the second I-O and paired pulse analysis. Conventional absolute and normalized baseline and post-tetanic EPSP slope and PS measures are presented in Supplemental data (Supplemental Material Figure S2).

#### Assessment of the synaptic contribution to PS plasticity in adult and aged male and female rats

To assess the contribution of synaptic input on enduring PS potentiation, absolute measures of the EPSP slope and PS were arranged in 30min bins and the EPSP slope measure of the first post-tetanic 30min bin (0-30min) was correlated (Pearson) with the three 30min post-tetanic PS measures (0-30min, 31-60, and 61-90min post-LTP). For contrast, the EPSP slope and PS measures are also presented from the 30min baseline period to illustrate pre-tetanus correlations.

#### Analysis of Indexed NMDA Receptor Current During Moderate Strength Tetanic LTP

To assess differences in NMDA receptor activation during tetanic in adult and aged adult male and female rats, total area under the curve (AUC) was measured first for each pulse within a 15 pulse stimulation train for each of the 20 burst stimulations, beginning at ∼10ms after the first pulse, a period determined previously to constitute NMDA receptor activation^11^. Total NMDA AUC was also summed across all 20 trains and also contrasted with the post-tetanic PS potentiation (see Supplemental Figure S3).

### Statistical Analysis

With the exception of correlative variables, data was analyzed using multifactor analysis of variance (baseline EPSP slope and PS) or mixed design analysis of variance (age, sex, variable). Fisher’s LSD tests were used for post-hoc assessments. All analyses were performed using Statistica (StatSoft).

## RESULTS

### Female rats show greater intrinsic granule cell excitability than males both pre and post LTP. In adult male rats the LTP protocol induces an increase in excitability but does not induce an excitability increase in female rats

The involvement of intrinsic excitability in MPP LTP in adult male rats is consistent with the findings of Lopez-Rojas^2^. The greater normalized LTP in adult male than adult female rats (*Supplemental Figure S2*) replicates Maren^12^. Higher levels of excitability in the MPP circuit of female rats than male rats are seen in data sets from earlier studies^13^ (raw data shared by Safari and Karimi 2021), but see ^14^ for counter example.

Adult female rats have a leftward shift in E-S coupling relative to same age males, however aged males and females have a similar E-S coupling EPSP slopes but with the distinction that at higher EPSP values, PS increases in females but reach a ceiling effect, while PS values in aged males are ∼50% of female PS values (Fig. 1). This argues that higher female intrinsic excitability, while possibly diminished, is still present.

### Increases in the population spike post-LTP in female rats depend on increases in EPSP slope

In female rats, the PS-LTP following tetani is predicted by the EPSP slope increase occurring in the first 30 min post-tetani (Fig.2). This post-tetani EPSP-spike correlation is not significant for male rats. However, aged males also fail to show significant normalized slope or spike increases following tetani. These outcomes argue that of the two MPP plasticity mechanisms identified by Bliss and Lomo^1^, an increase in synaptic size drives an increase in population spike in dentate gyrus of female rats, while in adult male rats plasticity depends on an increased in granule cell dendritic excitability as shown by Lopez^2^. All groups showed a decrease in spike latency following tetani (Supplementary Material Fig. S2). This outcome is consistent with Bliss and Lomo’s report that spike latency decrease was the most consistent response to LTP protocols^1^. In a new study of MPP EPSP slope potentiation, Amani et al^15^ report a decline in normalized EPSP potentiation beginning as early as 8 mo in male rats. The present failure of older males to exhibit EPSP potentiation corroborates their finding.

**Figure 2.**
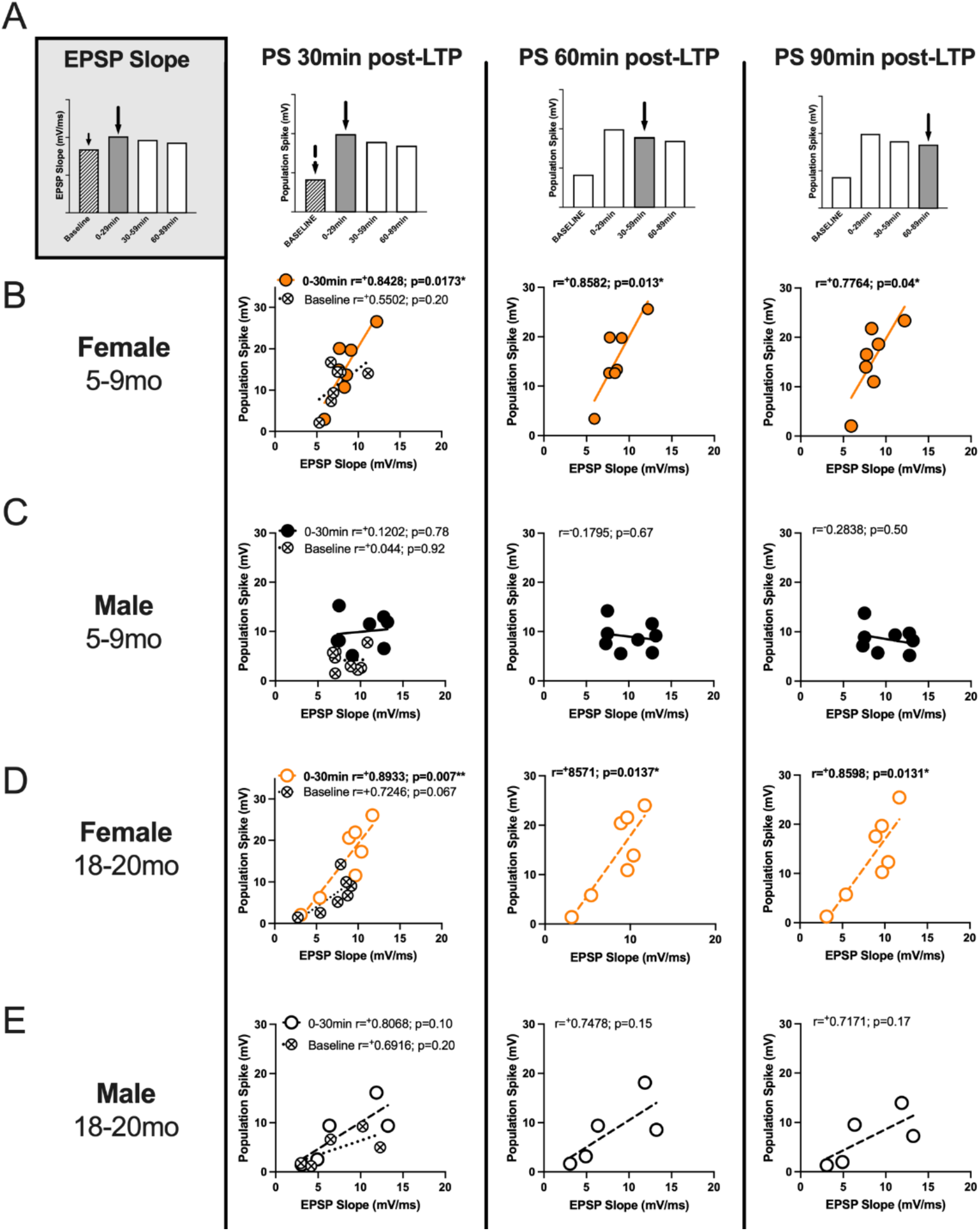
The effects of a moderate strength tetanic LTP protocol on the perforant path-dentate gyrus evoked population spike in young (5-7mo) and old (18-20mo) urethane anesthetized female rats, correlate with early EPSP slope potentiation. **A**. Absolute EPSP slope and PS values were organized into 30min bins (baseline, 0-30, 30-60, 60-90min post-LTP). EPSP slope values for the baseline period, and 0-30min post-LTP (early increases) were plotted against the baseline and the 0-90min PS binned values. Baseline correlations indicate that EPSP slope is not significantly correlated with PS values in any sex, or age group (crossed circles, B-E), however post-LTP EPSP slope values are predictive of increases in PS post-LTP in female rats (adult and aged; see bolded correlations in B and D). This effect was not observed in adult, or aged adult male rats (C and E).

### NMDA burst activation was stronger in adult males than in adult females or aged males

Using Maren’s approach^11^ of examining post-burst depolarization to evaluate NMDA activation by tetani, we found the three two three trains provided evidence of greater adult male (5-9mo) NMDA burst depolarization than that seen in female rats of the same age (Fig. 3). Consistent with this finding, Maren et al.^11^ reported larger NMDA burst depolarization in adult males than adult females. NMDA burst depolarization here decreased successive trains across the sexes, in both age groups. The outcome suggests NMDA induction is weakest in adult females, but no aged females.

**Figure 3.**
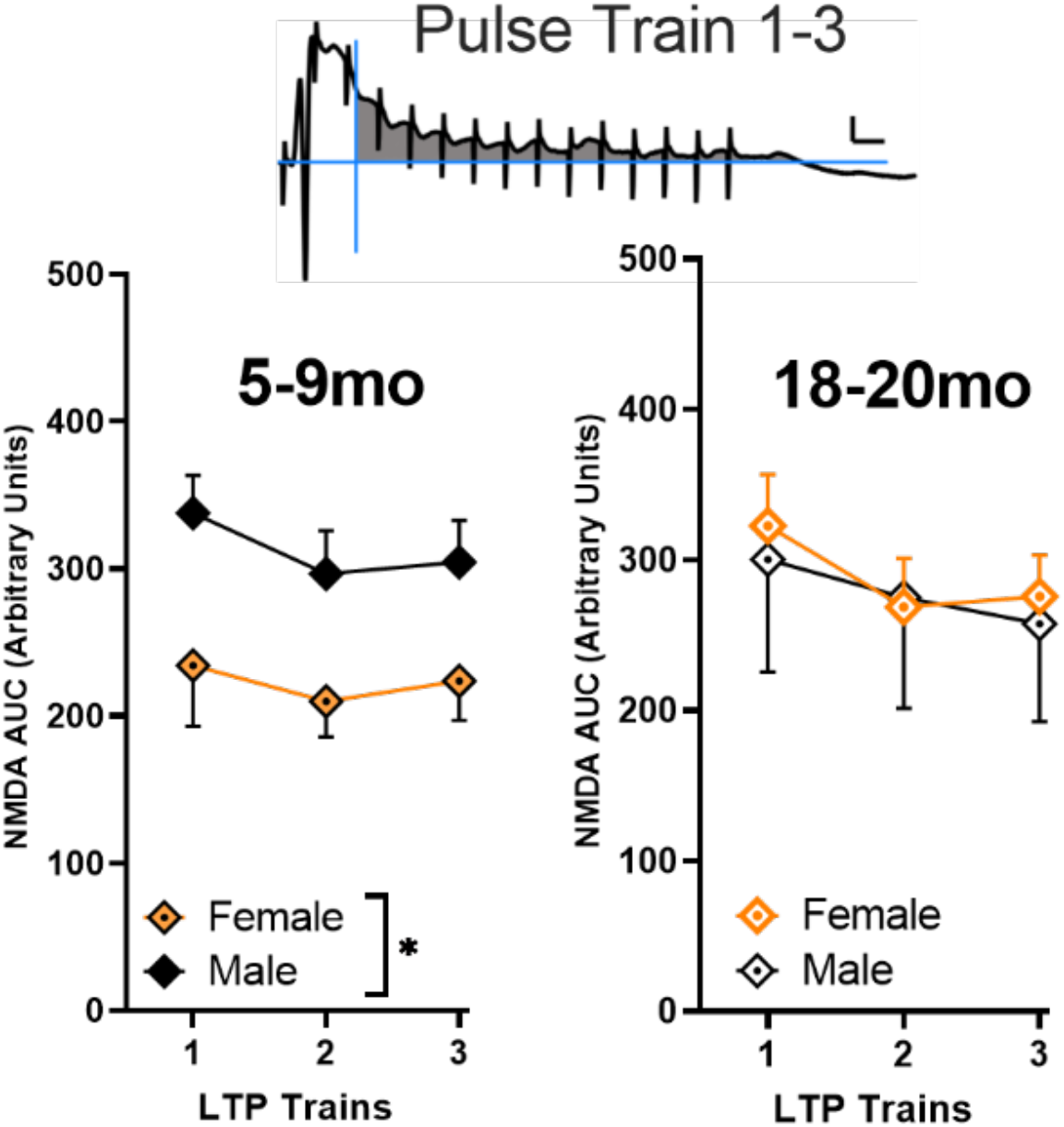
Assessment of the total NMDA receptor depolarization (Maren^11^) for the first 3 trains during tetanic stimulation. The male adult rats (5-9mo, left panel) had a larger NMDA-dependent component than adult female rats (F_1,13_=5.1936; p=0.04). In aged adult rats (right panel) there were no sex-dependent differences observed. In both age groups there was a significant effect of trial with successive trains decreasing in the total NMDA-dependent response to the tetanic stimulation rats (adult, F_2,26_=4.1925; p=0.02; aged adult, F_2,20_=11.567; p=0.0005). Scale is 4mV/5ms.

**Figure 4.**
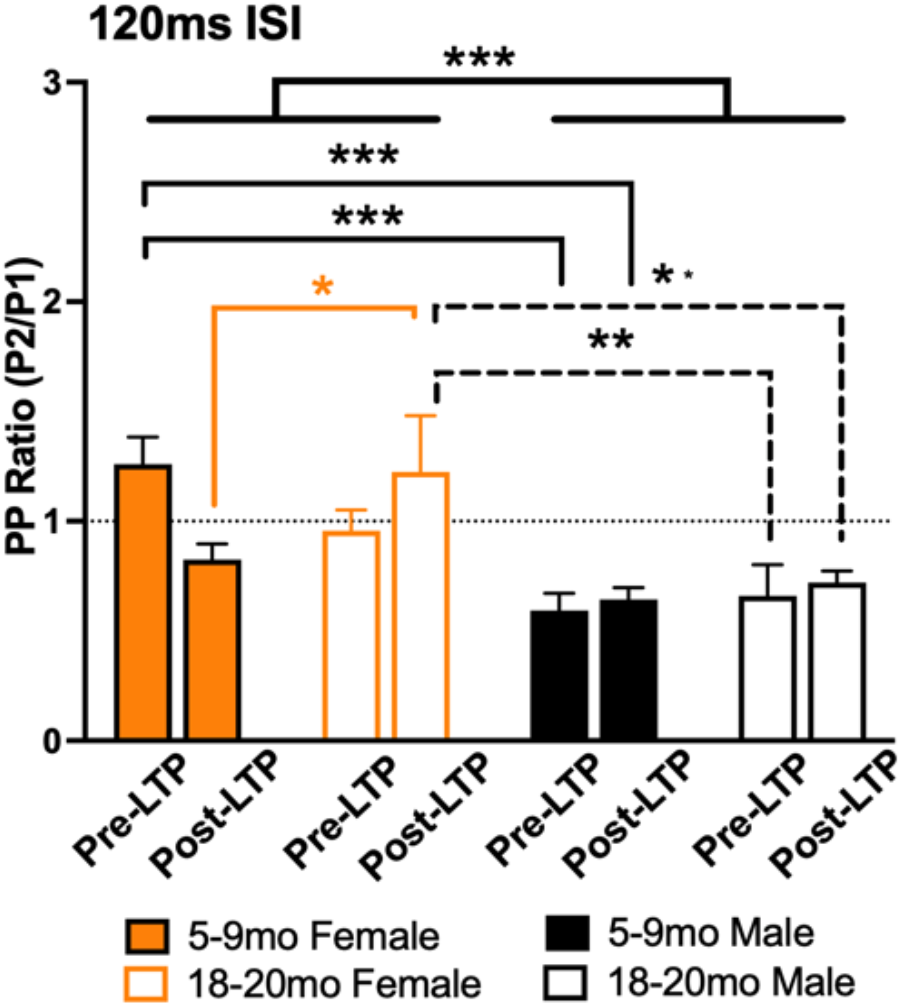
Paired pulse inhibition (120ms ISI) and facilitation of the PP-DG evoked population spike in young and old male and female rats. **A**. P2/P1 ratio for GABA-B dependent PPI (in 5-9mo and 18-20mo male and female rats at current levels used for baseline recordings; 50% maximal PS on P1), and same current Post-LTP. Female rats demonstrated less PPI compared to male rats. There was a significant age, sex, LTP interaction (F_1,23_)=4.3969; p<0.05. Data represent means± s.e.m. *=minimum p<0.05, **p<0.01 and ***p<0.001. **See Supplemental Figure 4 (S4) for full description of PP Input-Output profiles**.

### Paired pulse data revealed less GABA-B inhibitory modulation in female than male rats

Paired pulse inhibition probed at a 120 msec interval revealed the expected inhibition in male rats when the P1/P2 ratio was summed over all currents (∼50% inhibition). Females, however, did not show evidence of 120 msec ISI inhibition with their P2/P1 ratios averaging close to or above 1.0. Reduced GABA-B-mediated inhibition, indexed by this probe, may contribute to greater female intrinsic excitability. Canning and Leung^16^ demonstrated that in vivo granule cell excitability is controlled by GABA-B mediated inhibition. See Supplemental Figure S4 for full profile of Input-Output current curve paired pulse results.

## DISCUSSION

The present experiments provide evidence for both sex- and age-related differences in MPP plasticity in rat dentate gyrus. The sex difference in intrinsic excitability requires replication. While re-examining control data in earlier studies with other objectives provides some support for our findings, only one other laboratory has specifically addressed this variable. Maren’s studies did find greater normalized spike potentiation among Sprague-Dawley males under two kinds of anesthesia, but they specifically matched male and female rats for spike size potentially eliminating the differences in absolute spike/current relationships seen here.

What sex differences might account for these outcomes? Erwin and Cembrowski^17^ demonstrate that a subtype of mature granule cell, the PENK-expressing granule cell is more excitable than other subtypes and due to its greater excitability is preferentially recruited to spatial maps. They argue that this subtype dominates recruitment in hippocampally-dependent behaviors and supports sparse spatial representations in male mice. The critical role of intrinsic excitability in determining dentate gyrus MPP output was also recently highlighted by Zhang etal^18^. Oule et al^19^ identified the potassium channels responsible for dendritic intrinsic excitability in male mice as the Kv4.2 subtype and revealed that these channels are critical for spatial pattern separation.

In a new study identifying PENK-expressing granule cells in both male and female rats, Johnson et al^20^ demonstrate that female dorsal dentate gyrus contains a significantly larger proportion of PENK-expressing granule cells than male dentate gyrus. The higher PENK-cell proportion in females may support their greater intrinsic excitability. Based on the data from mice MPP-supported spatial and nonspatial representations might be predicted to be less sparse, but possibly more robust, in female rats than males (see for example O’Leary et al^21^, Olave et al^22^ Zhvania et al^23^). Experimental evidence with respect to place field and episodic representation in female versus male rats is needed. In an evolutionary context^24^ spatial map demands appear different for male and female rats. *Rattus norwegicus* has spatial territories an order of magnitude larger than females^25^ and may require better spatial resolution for those territories.

The role of aging in dentate gyrus plasticity has been more extensively studied. A selective decrease in MPP synapses on granule cells dendrites (3 mo versus 28 mo) was reported in 1976^26^. Barnes ^27^ (comparing behavioral and LTP measures in 10-16 mo versus 28-34 mo old male rats found repeated trains produced enduring EPSP slope potentiation at 10-16 mo but a declining potentiation in the senescent rats. Barnes et al^28^ report reduced NMDA currents and a higher threshold for EPSP LTP in senescent rats, which were memory-impaired (see also Yang et al^29^). Their middle-aged group (9-10 mo), are similar to the adult rats here and appeared intermediate between young and senescent rats in depolarization needed for EPSP slope potentiation. In the Barnes et al. study, the NMDA currents in female rats were not examined and warrant further investigation. Arc-active granule cells with spatial exploration decline across young, middle and aged rats^30^.

Multiple laboratories^31, 32^ have demonstrated both memory-impaired and memory-unimpaired aged male rats are seen when probed on hippocampally-dependent tasks, thus variability in initiating and maintaining plasticity with age is likely. Understanding the mechanistic underpinning of that variability will be useful for cognitive anti-aging strategies. Lubec et al^32^ working with Sprague-Dawley rats aged 22-24 mo and comparing them to 6 mo rats identified proteomic changes that occurred differentially in aged impaired and unimpaired rats. Impaired rats were deficient in pathways related to energy metabolism and potassium ion regulation. They found unimpaired rats differed from the general population as early as 6 mo of age corroborating the Amani et al^15^ assertion that decreases in MPP plasticity are an early harbinger of aging. Lubec et al^33^ later found that increasing dopamine in aged male Sprague-Dawley rats with both intermediate and severe behavioral impairments on hippocampally-dependent tasks restored behavior to that of young rats and restored spine numbers on granule cells to young levels. Both spatial behavior and cognitive flexibility were improved.

The present experiments suggest MPP-related aging changes are likely to differ among males and females with males being more vulnerable to aging-related disruption of plasticity and likely to show greater impairment on hippocampally-dependent tasks. This hypothesis remains to be explored.

## FUNDING ACKNOWLEDGEMENTS

This work was supported by Natural Science and Engineering Research Council of Canada (Discovery) and Alzheimer Society of Canada to SGW.

## CONFLICT OF INTEREST STATEMENT

The authors declare we have no conflict of interest.

## Notes

### Competing Interest Statement

The authors have declared no competing interest.

## REFERENCES

(1) Bliss, T. V.; Lomo, T. Long-lasting potentiation of synaptic transmission in the dentate area of the anaesthetized rabbit following stimulation of the perforant path. The Journal of physiology 1973, 232 (2), 331–356. DOI: 10.1113/jphysiol.1973.sp010273

(2) Lopez-Rojas, J.; Heine, M.; Kreutz, M. R. Plasticity of intrinsic excitability in mature granule cells of the dentate gyrus. Scientific reports 2016, 6, 21615. DOI: 10.1038/srep21615

(3) Daoudal, G.; Debanne, D. Long-term plasticity of intrinsic excitability: learning rules and mechanisms. Learning & memory 2003, 10 (6), 456–465. DOI: 10.1101/lm.64103

(4) Hubert, M. F.; Laroque, P.; Gillet, J. P.; Keenan, K. P. The effects of diet, ad Libitum feeding, and moderate and severe dietary restriction on body weight, survival, clinical pathology parameters, and cause of death in control Sprague-Dawley rats. Toxicol Sci 2000, 58 (1), 195–207. DOI: 10.1093/toxsci/58.1.195

(5) Davis, S.; Salin, H.; Helme-Guizon, A.; Dumas, S.; Stephan, A.; Corbex, M.; Mallet, J.; Laroche, S. Dysfunctional regulation of alphaCaMKII and syntaxin 1B transcription after induction of LTP in the aged rat. The European journal of neuroscience 2000, 12 (9), 3276–3282. DOI: 10.1046/j.1460-9568.2000.00193.x

(6) Joy, R. M.; Albertson, T. E. NMDA receptors have a dominant role in population spike-paired pulse facilitation in the dentate gyrus of urethane-anesthetized rats. Brain research 1993, 604 (1-2), 273–282. Ruan, D. Y.; Chen, J. T.; Zhao, C.; Xu, Y. Z.; Wang, M.; Zhao, W. F. Impairment of long-term potentiation and paired-pulse facilitation in rat hippocampal dentate gyrus following developmental lead exposure in vivo. Brain research 1998, 806 (2), 196–201. DOI: 10.1016/s0006-8993(98)00739-2

(7) Brucato, F. H.; Morrisett, R. A.; Wilson, W. A.; Swartzwelder, H. S. The GABAB receptor antagonist, CGP-35348, inhibits paired-pulse disinhibition in the rat dentate gyrus in vivo. Brain research 1992, 588 (1), 150–153. DOI: 10.1016/0006-8993(92)91355-i

(8) Walling, S. G.; Harley, C. W. Locus ceruleus activation initiates delayed synaptic potentiation of perforant path input to the dentate gyrus in awake rats: a novel beta-adrenergic-and protein synthesis-dependent mammalian plasticity mechanism. The Journal of neuroscience : the official journal of the Society for Neuroscience 2004, 24 (3), 598–604. DOI: 10.1523/JNEUROSCI.4426-03.2004.

(9) Straube, T.; Frey, J. U. Involvement of β-adrenergic receptors in protein synthesis-dependent late long-term potentiation (LTP) in the dentate gyrus of freely moving rats: the critical role of the LTP induction strength. Neuroscience 2003, 119 (2), 473–479. DOI: 10.1016/s0306-4522(03)00151-9.

(10) Straube, T.; Frey, J. U. Involvement of beta-adrenergic receptors in protein synthesis-dependent late long-term potentiation (LTP) in the dentate gyrus of freely moving rats: the critical role of the LTP induction strength. Neuroscience 2003, 119 (2), 473–479. DOI: 10.1016/s0306-4522(03)00151-9

(11) Maren, S.; Baudry, M.; Thompson, R. F. Effects of the novel NMDA receptor antagonist, CGP 39551, on field potentials and the induction and expression of LTP in the dentate gyrus in vivo. Synapse 1992, 11 (3), 221–228. DOI: 10.1002/syn.890110307

(12) Maren, S.; De Oca, B.; Fanselow, M. S. Sex differences in hippocampal long-term potentiation (LTP) and Pavlovian fear conditioning in rats: positive correlation between LTP and contextual learning. Brain research 1994, 661 (1-2), 25–34. DOI: 10.1016/0006-8993(94)91176-2 From NLM Medline. Maren, S. Sexually dimorphic perforant path long-term potentiation (LTP) in urethane-anesthetized rats. Neurosci Lett 1995, 196 (3), 177–180.

(13) Kehoe, P.; Bronzino, J. D. Neonatal stress alters LTP in freely moving male and female adult rats. Hippocampus 1999, 9 (6), 651–658. DOI: 10.1002/(SICI)1098-1063(1999)9:6<651::AID-HIPO6>3.0.CO;2-P From NLM Medline. Zitman, F. M.; Richter-Levin, G. Age and sex-dependent differences in activity, plasticity and response to stress in the dentate gyrus. Neuroscience 2013, 249, 21–30. DOI: 10.1016/j.neuroscience.2013.05.030 Safari, S.; Ahmadi, N.; Mohammadkhani, R.; Ghahremani, R.; Khajvand-Abedeni, M.; Shahidi, S.; Komaki, A.; Salehi, I.; Karimi, S. A. Sex differences in spatial learning and memory and hippocampal long-term potentiation at perforant pathway-dentate gyrus (PP-DG) synapses in Wistar rats. Behav Brain Funct 2021, 17 (1), 9. DOI: 10.1186/s12993-021-00184-y

(14) Elmarzouki, H.; Aboussaleh, Y.; Bitiktas, S.; Suer, C.; Artis, A. S.; Dolu, N.; Ahami, A. Effects of cold exposure on behavioral and electrophysiological parameters related with hippocampal function in rats. Frontiers in cellular neuroscience 2014, 8, 253. DOI: 10.3389/fncel.2014.00253

(15) Amani, M.; Lauterborn, J. C.; Le, A. A.; Cox, B. M.; Wang, W.; Quintanilla, J.; Cox, C. D.; Gall, C. M.; Lynch, G. Rapid Aging in the Perforant Path Projections to the Rodent Dentate Gyrus. The Journal of neuroscience : the official journal of the Society for Neuroscience 2021, 41 (10), 2301–2312. DOI: 10.1523/JNEUROSCI.2376-20.2021

(16) Canning, K. J.; Leung, L. S. Excitability of rat dentate gyrus granule cells in vivo is controlled by tonic and evoked GABA(B) receptor-mediated inhibition. Brain research 2000, 863 (1-2), 271–275. DOI: 10.1016/s0006-8993(00)02124-7

(17) Erwin, S. R.; Sun, W.; Copeland, M.; Lindo, S.; Spruston, N.; Cembrowski, M. S. A Sparse, Spatially Biased Subtype of Mature Granule Cell Dominates Recruitment in Hippocampal-Associated Behaviors. Cell reports 2020, 31 (4), 107551. DOI: 10.1016/j.celrep.2020.107551

(18) Zhang, X.; Schlogl, A.; Jonas, P. Selective Routing of Spatial Information Flow from Input to Output in Hippocampal Granule Cells. Neuron 2020, 107 (6), 1212–1225 e1217. DOI: 10.1016/j.neuron.2020.07.006

(19) Oule, M.; Atucha, E.; Wells, T. M.; Macharadze, T.; Sauvage, M. M.; Kreutz, M. R.; Lopez-Rojas, J. Dendritic Kv4.2 potassium channels selectively mediate spatial pattern separation in the dentate gyrus. iScience 2021, 24 (8), 102876. DOI: 10.1016/j.isci.2021.102876

(20) Johnson, M. A.; Contoreggi, N. H.; Kogan, J. F.; Bryson, M.; Rubin, B. R.; Gray, J. D.; Kreek, M. J.; McEwen, B. S.; Milner, T. A. Chronic stress differentially alters mRNA expression of opioid peptides and receptors in the dorsal hippocampus of female and male rats. The Journal of comparative neurology 2021, 529 (10), 2636–2657. DOI: 10.1002/cne.25115

(21) O’Leary, T. P.; Askari, B.; Lee, B. H.; Darby, K.; Knudson, C.; Ash, A. M.; Seib, D. R.; Espinueva, D. F.; Snyder, J. S. Sex Differences in the Spatial Behavior Functions of Adult-Born Neurons in Rats. eNeuro 2022, 9 (3). DOI: 10.1523/ENEURO.0054-22.2022

(22) Olave, F. A.; Aguayo, F. I.; Roman-Albasini, L.; Corrales, W. A.; Silva, J. P.; Gonzalez, P. I.; Lagos, S.; Garcia, M. A.; Alarcon-Mardones, M.; Rojas, P. S.; et al. Chronic restraint stress produces sex-specific behavioral and molecular outcomes in the dorsal and ventral rat hippocampus. Neurobiol Stress 2022, 17, 100440. DOI: 10.1016/j.ynstr.2022.100440

(23) Zhvania, M. G.; Japaridze, N.; Tizabi, Y.; Lomidze, N.; Pochkhidze, N.; Lordkipanidze, T. Age-related cognitive decline in rats is sex and context dependent. Neurosci Lett 2021, 765, 136262. DOI: 10.1016/j.neulet.2021.136262

(24) Sherry, D. F.; Jacobs, L. F.; Gaulin, S. J. Spatial memory and adaptive specialization of the hippocampus. Trends in neurosciences 1992, 15 (8), 298–303. DOI: 10.1016/0166-2236(92)90080-r

(25) Oyedele, D. T.; Sah, S. A.; Kairuddinand, L.; Wan Ibrahim, W. M. Range Measurement and a Habitat Suitability Map for the Norway Rat in a Highly Developed Urban Environment. Trop Life Sci Res 2015, 26 (2), 27–44.

(26) Bondareff, W.; Geinisman, Y. Loss of synapses in the dentate gyrus of the senescent rat. Am J Anat 1976, 145 (1), 129–136. DOI: 10.1002/aja.1001450110

(27) Barnes, C. A. Memory deficits associated with senescence: a neurophysiological and behavioral study in the rat. Journal of comparative and physiological psychology 1979, 93 (1), 74–104. DOI: 10.1037/h0077579

(28) Barnes, C. A.; Rao, G.; Houston, F. P. LTP induction threshold change in old rats at the perforant path--granule cell synapse. Neurobiology of aging 2000, 21 (5), 613–620. DOI: 10.1016/s0197-4580(00)00163-9

(29) Yang, Z.; Krause, M.; Rao, G.; McNaughton, B. L.; Barnes, C. A. Synaptic commitment: developmentally regulated reciprocal changes in hippocampal granule cell NMDA and AMPA receptors over the lifespan. Journal of neurophysiology 2008, 99 (6), 2760–2768. DOI: 10.1152/jn.01276.2007

(30) Small, S. A.; Chawla, M. K.; Buonocore, M.; Rapp, P. R.; Barnes, C. A. Imaging correlates of brain function in monkeys and rats isolates a hippocampal subregion differentially vulnerable to aging. Proceedings of the National Academy of Sciences of the United States of America 2004, 101 (18), 7181–7186. DOI: 10.1073/pnas.0400285101.

(31) Rapp, P. R.; Gallagher, M. Preserved neuron number in the hippocampus of aged rats with spatial learning deficits. Proceedings of the National Academy of Sciences of the United States of America 1996, 93 (18), 9926–9930. DOI: 10.1073/pnas.93.18.9926

(32) Lubec, J.; Smidak, R.; Malikovic, J.; Feyissa, D. D.; Korz, V.; Hoger, H.; Lubec, G. Dentate Gyrus Peroxiredoxin 6 Levels Discriminate Aged Unimpaired From Impaired Rats in a Spatial Memory Task. Front Aging Neurosci 2019, 11, 198. DOI: 10.3389/fnagi.2019.00198

(33) Lubec, J.; Kalaba, P.; Hussein, A. M.; Feyissa, D. D.; Kotob, M. H.; Mahmmoud, R. R.; Wieder, O.; Garon, A.; Sagheddu, C.; Ilic, M.; et al. Reinstatement of synaptic plasticity in the aging brain through specific dopamine transporter inhibition. Molecular psychiatry 2021, 26 (12), 7076–7090. DOI: 10.1038/s41380-021-01214-x

