## Supplemental Material Figures S1-S4 for "Sprague-Dawley rats differ in responses to medial perforant path paired pulse and tetanic activation as a function of sex and age"

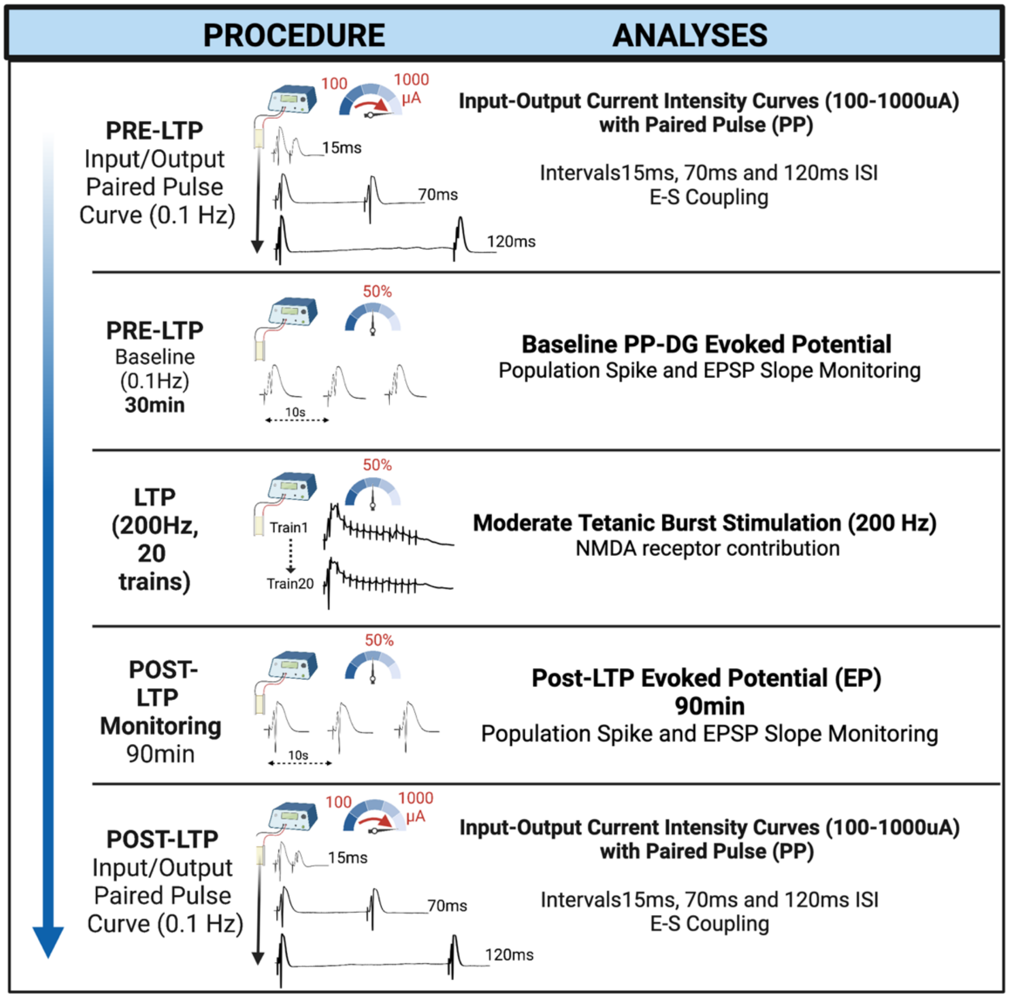


***Supplemental Figure S1***

Graphical experimental procedure. In brief, experiment commenced with paired pulse (15, 70, and 120ms interleaved on a current intensity profile (100-1000µA). Baseline recording (min 30min) with current eliciting 50% maximal PS. Moderate strength tetanic stimulation (Protocol B, Straube and Frey[9]), followed by 90min recording. Post-LTP PP and input-output current intensity curve.

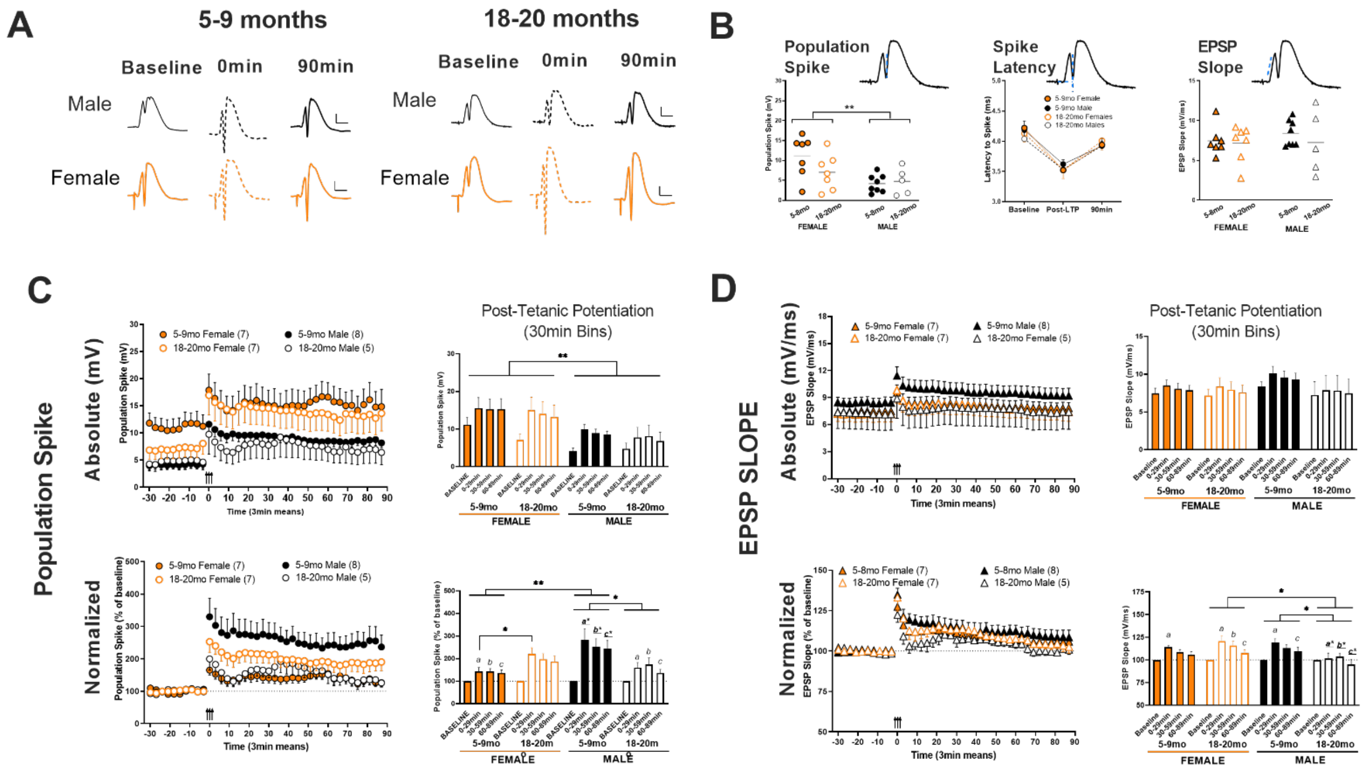


*Supplemental Figure S2.*

**The effects of a moderate strength tetanic LTP protocol on the perforant path- dentate gyrus evoked population spike (PS) and EPSP slope measures in adult (5-9mo) and aged adult (18-20mo) urethane anesthetized male and female rats**. **A.** Sample waveforms for three periods before (baseline) and after (0 and 90min) moderate strength tetanic stimulation. Scale is 4mV and 5ms. **B.** Absolute PS (baseline current), PS latency (baseline, 0-, and 90min post LTP) and EPSP slope measures (baseline current). PS values (mV) were significantly higher in female rats compared to male rats at baseline current levels, but EPSP slope (mV/ms) were not different. **C.** Temporal profile (X-Y plot), and 30min binned data (bar graph) of absolute (top panels), and normalized (bottom panels) PS data. Female rats had significantly larger PS (mV) amplitudes than male rats (main effect sex, $F_{1,23}$=7.378; p=0.012). Normalization of PS data illustrates adult males (5-9mo) had higher percentage PS increases than aged males (18-20mo), and adult female rats (age, sex, LTP interaction, $F_{3,69}$=6.52; p=0.0006, with post-hoc). **D.** EPSP Slope data. No sex- or age-dependent differences were observed in absolute EPSP slope measures (top panels). When data was normalized to baseline averages however a significant age, sex, slope interaction was revealed ($F_{3,69}$=3.660; p=0.016, with post-hoc). Aged male rats had lower EPSP slope percentage increases post-LTP compared to younger adult and aged female rats. Normalized baseline measures were not included in these analyses in C or D. *=minimum p<0.05, **p<0.01. Lower case letters are different from same underlined* letters in an examination of binned intervals e.g. *b different from* *b**.

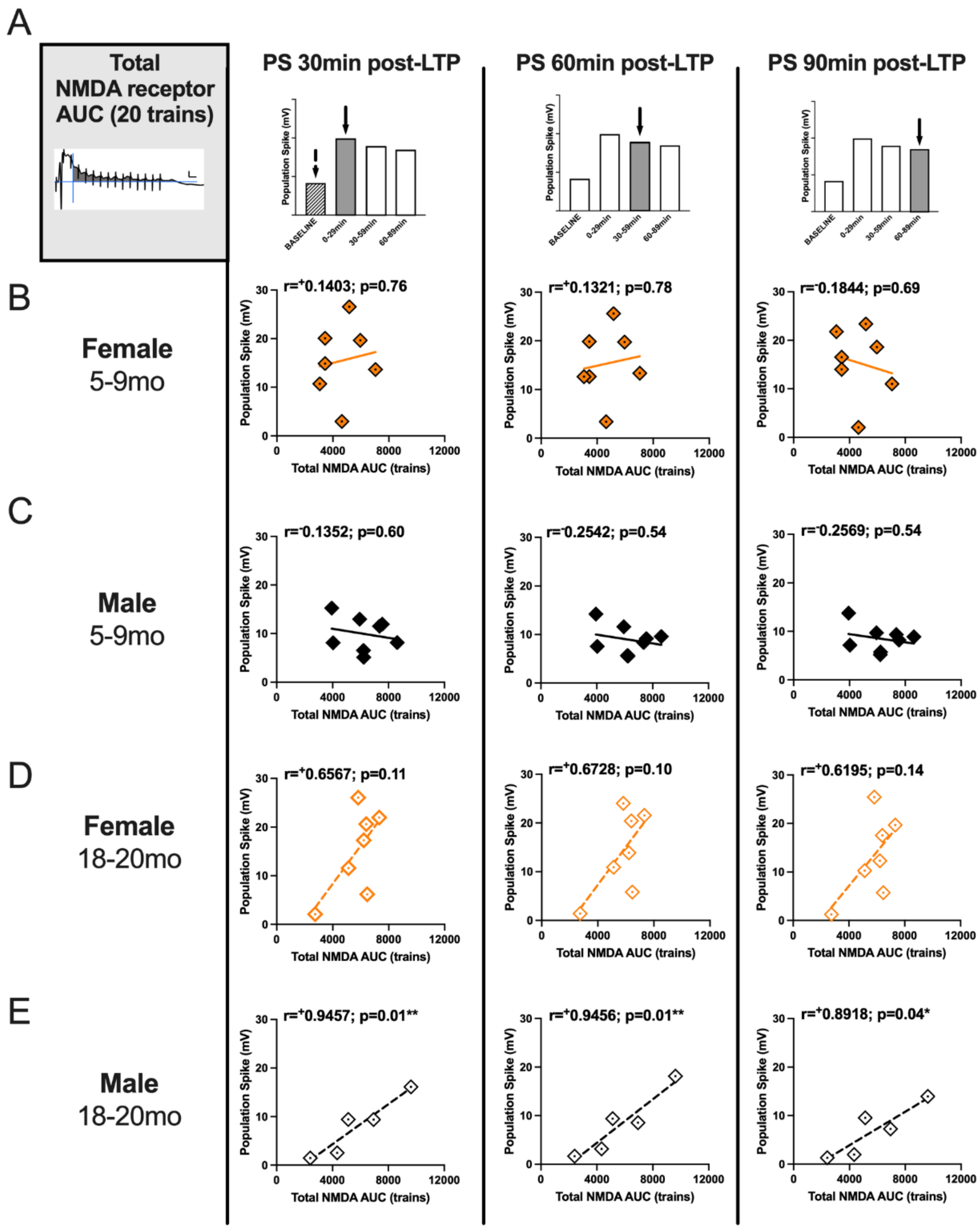


***Supplemental Figure S3.*** **Total NMDA receptor contribution during moderate tetanic LTP stimulation correlated with population spike increase in 30min bins in adult and aged male and female rats.** The total NMDA receptor Area Under the Curve (AUC) for the 15 pulse, and 20 tetanic trains was plotted against the absolute PS amplitude (mV) for the post-tetanic period (0-90min post-LTP). Total NMDA AUC was not correlated with absolute PS values (A-B) however, a significant correlation was more associated with aged male (18-20mo, see E), and trend in aged female rats (D).

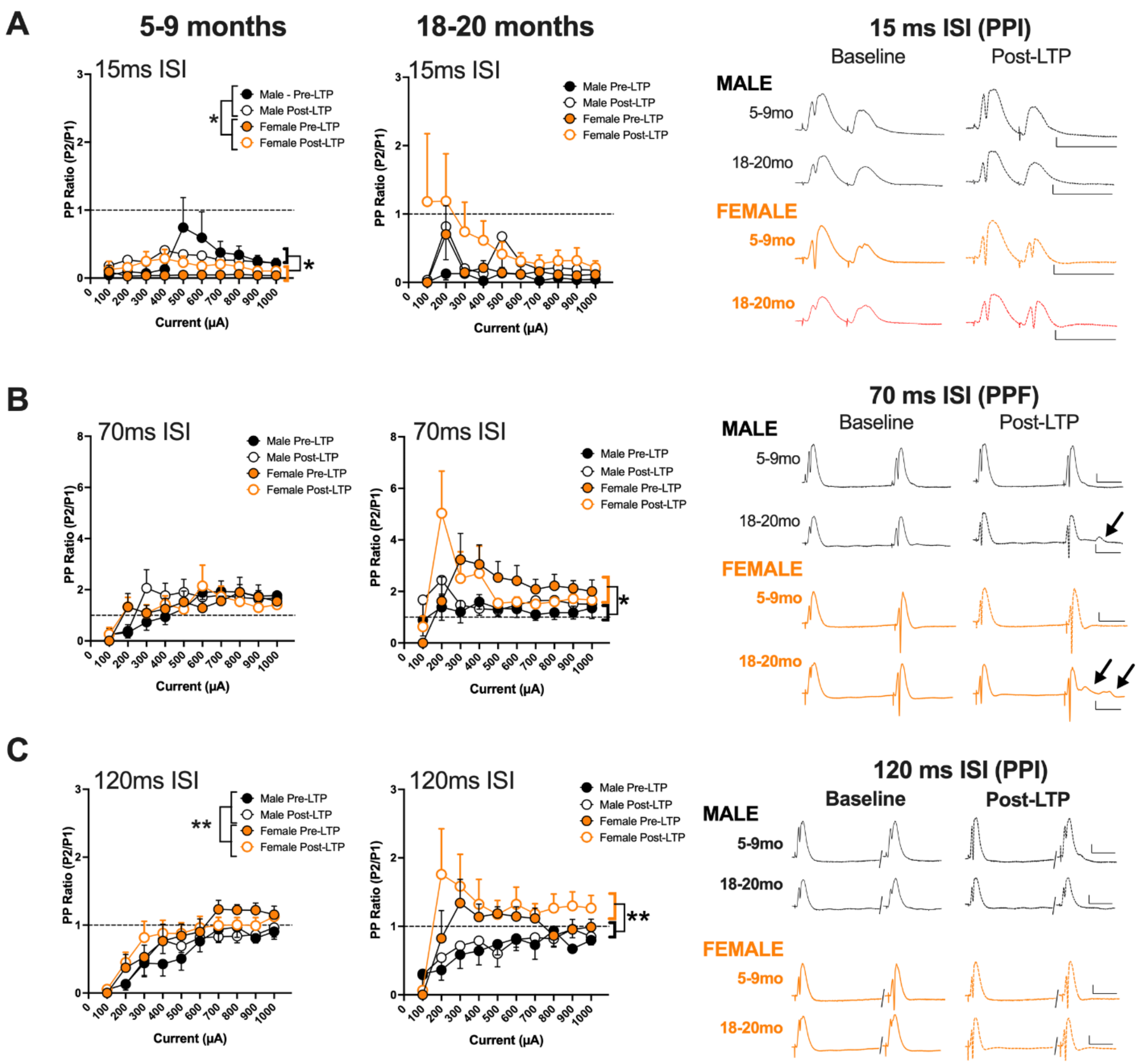


***Supplemental Figure S4.***

**The effects of a moderate tetanic LTP protocol on paired pulse ratio input-output current intensity curves for male and female adult and aged adult Sprague Dawley rats at three interstimulus intervals. A.** 15ms ISI (PPI); no significant main effects or interactions of sex for either age groups were observed. **B.** 70ms ISI (PPF); no significant main effects or interactions of sex for either age groups, however examples of hyperexcitability emerged in some male and female rats (examples shown, arrows in waveforms). These could not be quantified. **C.** 120ms ISI (PPI): In the 5-9mo age group there was a significant main effect of Age ($F_{1,13}$=18.46; p<0.001), and in the 18-20mo rats, a significant current x sex interaction ($F_{9,90}$=3.106; p=0.003). Data represent means± s.e.m. **p<0.01 and ***p<0.001. Scale in waveforms is 4mV/5ms.
